# FAM83G/PAWS1 controls cytoskeletal dynamics and cell migration through association with the SH3 adaptor CD2AP

**DOI:** 10.1101/106971

**Authors:** Timothy D. Cummins, Kevin Z. L. Wu, Polyxeni Bozatzi, Kevin S. Dingwell, Thomas Macartney, Nicola T Wood, Joby Varghese, Robert Gourlay, David G Campbell, Alan Prescott, Eric Griffis, James C Smith, Gopal P Sapkota

## Abstract

Our previous studies of PAWS1 (**P**rotein **A**ssociated **W**ith **S**MAD**1**) have suggested that this molecule has roles beyond BMP signalling. To investigate these roles, we have used CRISPR/Cas9 to generate PAWS1 knockout cells. Here, we show that PAWS1 plays a role in the regulation of the cytoskeletal machinery, including actin and focal adhesion dynamics, and cell migration. Confocal microscopy and live cell imaging of actin in U2OS cells indicate that PAWS1 is also involved in cytoskeletal dynamics and organization. Loss of PAWS1 causes severe defects in F-actin organization and distribution as well as in lamellipodial organization, resulting in impaired cell migration. PAWS1 interacts in a dynamic fashion with the actin/cytoskeletal regulator CD2AP at lamellae, suggesting that its association with CD2AP controls actin organization and cellular migration.

**Summary statement:** PAWS1/FAM83G controls cell migration by influencing the organisation of F-actin and focal adhesions and the distribution of the actin stress fibre network through its association with CD2AP.

## Introduction

Cell migration is involved in embryonic development, wound healing, the immune response and cancer metastasis(Easley et al., 2008; Fife et al., 2014). Although many of the molecules and biophysical processes involved in cell migration have been identified and characterized (Huber et al., 2015; Leduc and Etienne-Manneville, 2015; Mohapatra et al., 2016), we do not have a complete understanding of the process. One of the most important properties of cell migration is the ability of cells to fine-tune their cytoskeletal structure in response to changing environmental cues such as growth factor stimulation (Dang et al., 2013; Krause and Gautreau, 2014; Mendoza et al., 2015; Timpson et al., 2011).

Cytoskeletal components such as actin and tubulin play important roles in migration and invasion, notably in the pathology of tumour cells(Shortrede et al., 2016). Actin takes two forms: monomeric globular (G-actin) and filamentous (F-actin). F-actin polymerization is responsible for dynamic changes in cell shape and for chemotactic responses to growth factor signalling. It is also involved in the formation of lamellipodia, filopodia and other macromembrane structures that drive directional or chemotactic migration(Johnson et al., 2015; Kelley et al., 2010; King et al., 2016; Welf et al., 2012). Without properly regulated actin polymerization and branching, cells are unable to properly sense their microenvironment and they may display unregulated migratory behaviour.

The organization and polymerization of actin are controlled by molecular complexes that include Arp2/3 (actin related protein 2/3) and WASP/WAVE regulators that are downstream of the small GTPases Rho, Rac and Cdc42 (Devreotes and Horwitz, 2015; Guo et al., 2006). Dynamic membrane structures such as invadopodia, lamellipodia and pseudopodia are formed through the regulation of actin polymerization by association with nucleators, crosslinkers, capping proteins, severing proteins, debranching proteins, and myosin motors(Chi et al., 2014; Lehtimaki et al., 2016; Mierke, 2015). One such regulator is the adaptor protein CD2AP, which delivers capping proteins to the barbed ends of polymerizing F-actin. The resulting change in network architecture leads to plasma membrane ruffling, chemotactic arching and eventually motility(Bruck et al., 2006; Tang and Brieher, 2013; Zhao et al., 2013). The branch-promoting activity of CD2AP, together with the capping proteins CAPZA1/B1, leads to modifications in branched actin and causes membrane distortion and changes in tight junctions (Tang and Brieher, 2013; Zhao et al., 2013).

PAWS1 is a member of the FAM83 family of proteins that is characterized by the presence of a conserved *DUF1669* domain of unknown function. The domain includes a pseudo-phospholipase D (PLD) catalytic motif, so-called because no PLD activity has been detected in FAM83 proteins (Cipriano et al., 2012; Cipriano et al., 2013). Outside the *DUF1669* domain, the FAM83 members are distinct, perhaps pointing to different roles for each member. We have previously shown that PAWS1 interacts with SMAD1 and modulates BMP signalling and transcription(Vogt et al., 2014); here we demonstrate that loss of PAWS1 causes profound morphological and migratory changes in cells. A proteomic screen of the FAM83 family of proteins reveals that in addition to the SMADs, PAWS1 interacts with CD2AP. Bearing in mind the roles of CD2AP in cytoskeletal organization, dynamics and cell migration, this observation suggests that PAWS1 might interact with CD2AP to regulate cytoskeletal machinery and cell migration(Bruck et al., 2006; Tang and Brieher, 2013; Zhao et al., 2013). Our results indicate that PAWS1 is a novel regulator of actin-cytoskeletal dynamics, cell locomotion and migration.

## Results

### PAWS1 deficiency affects cell morphology, cytoskeletal dynamics and migration

To investigate the functions of PAWS1, we generated PAWS1 knockout U2OS cells (PAWS1^-/-^) by CRISPR/Cas9 targeting of exon 2 of the PAWS1 gene (Figure 1A). The loss of PAWS1 protein in the isolated clone of U2OS cells was verified by western blotting (Figure 1B), while genomic sequencing surrounding the *sgRNA* target site revealed a 5-base pair deletion from both alleles (Figure 1A). For rescue experiments, we employed a previously described retroviral method(Vogt et al., 2014) to stably restore the expression of wild type PAWS1 in PAWS1^-/-^ cells (PAWS1^Res^). We note that levels of PAWS1 in PAWS1^Res^ cells were slightly higher than endogenous levels in control U2OS and HaCaT keratinocyte cells (Figure 1B). Under these conditions, bright-field microscopy revealed that compared with both wild type and PAWS1^Res^ cells, PAWS1^-/-^ U2OS cells displayed reduced lamellipodia, abnormal membrane structures, and delayed adhesion to plastic (Figure 1C). Consistent with the differences in cell morphology, phalloidin staining of fixed PAWS1^-/-^ U2OS cells showed a disorganized and tangled mesh of actin, while wild type U2OS cells and PAWS1^Res^ cells showed normal actin stress fibre organization (Figure 1C; Supplementary Figure 2A-C). Inspection of actin fibre organization in PAWS1^-/-^ and wild type U2OS cells revealed more filopodia-like protrusions in PAWS1^-/-^ cells compared with wild type cells (Supplementary Figure 2D&E).

**Figure 1.**
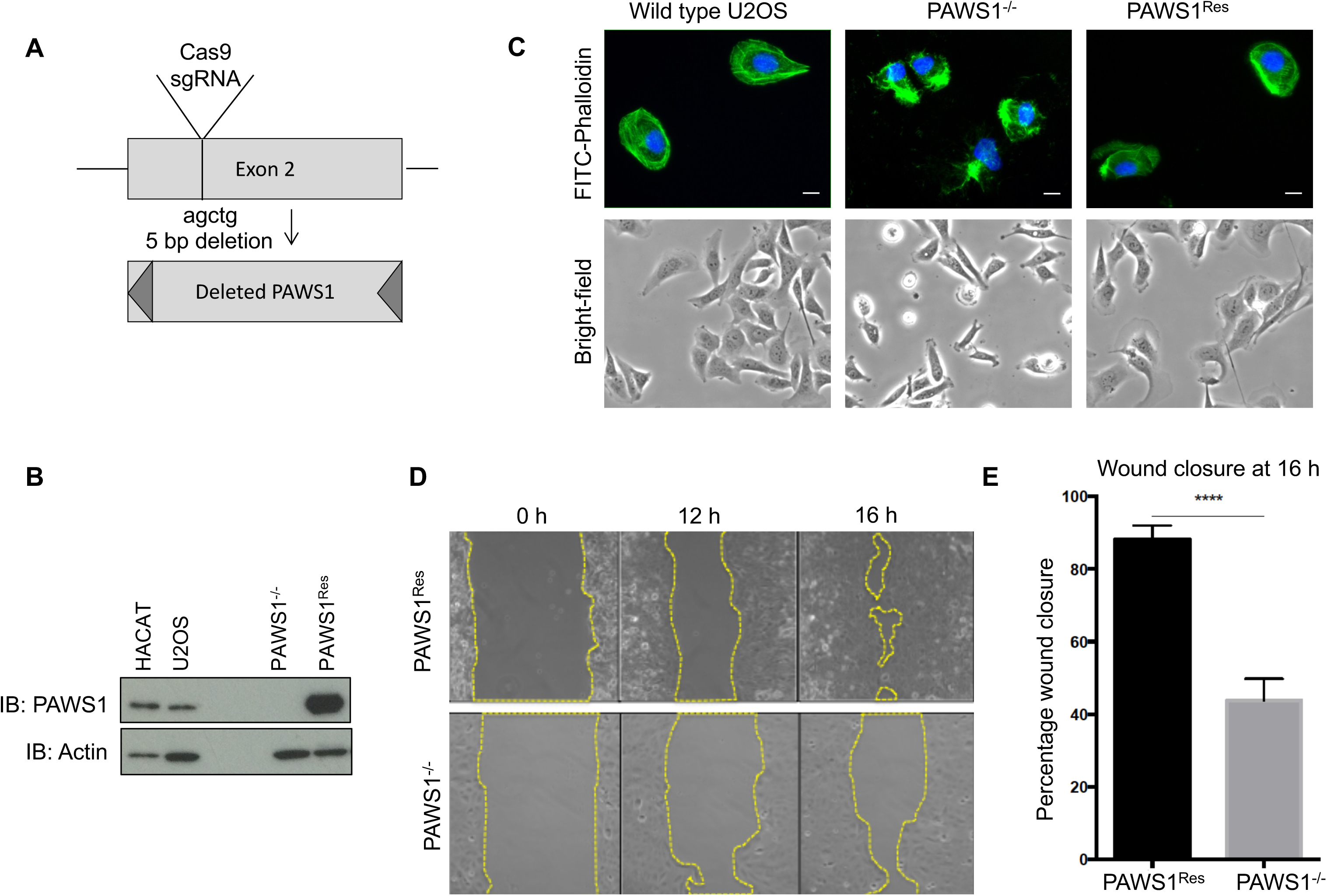
Loss of PAWS1 causes defects in U2OS cell migration and morphology. **A.** CRISPR mediated deletion of PAWS1 at exon 2 of the PAWS1 gene. **B.** Anti-PAWS1 immunoblots of 20 µg extracts from control HaCaT keratinocytes and U2OS osteosarcoma cells, as well as targeted PAWS1 knockout (PAWS1^-/-^) U2OS cells and knockout cells rescued with wild type PAWS1 (PAWS1^Res^). **C.** Fluorescence microscopy of actin (FITC-Phalloidin (green)) and DAPI (blue) staining in control U2OS cells, PAWS1^-/-^ cells or PAWS1^Res^ cells (Scale bar is 10 µm) together with bright-field microscopic images depicting actin organization and morphology in the indicated cell types. **D.** Time-lapse wound healing migration of PAWS1^-/-^ and PAWS1^Res^ U2OS cells at 0, 12 h and 16 h following removal of the insert separating wells of confluent cells. Images were taken using phase microscopy at 20X magnification. **E.** The percentage of wound (gap) closure at 16 h (as indicated in C) was quantified and plotted as shown (mean±SD; N=3).

**Figure 2.**
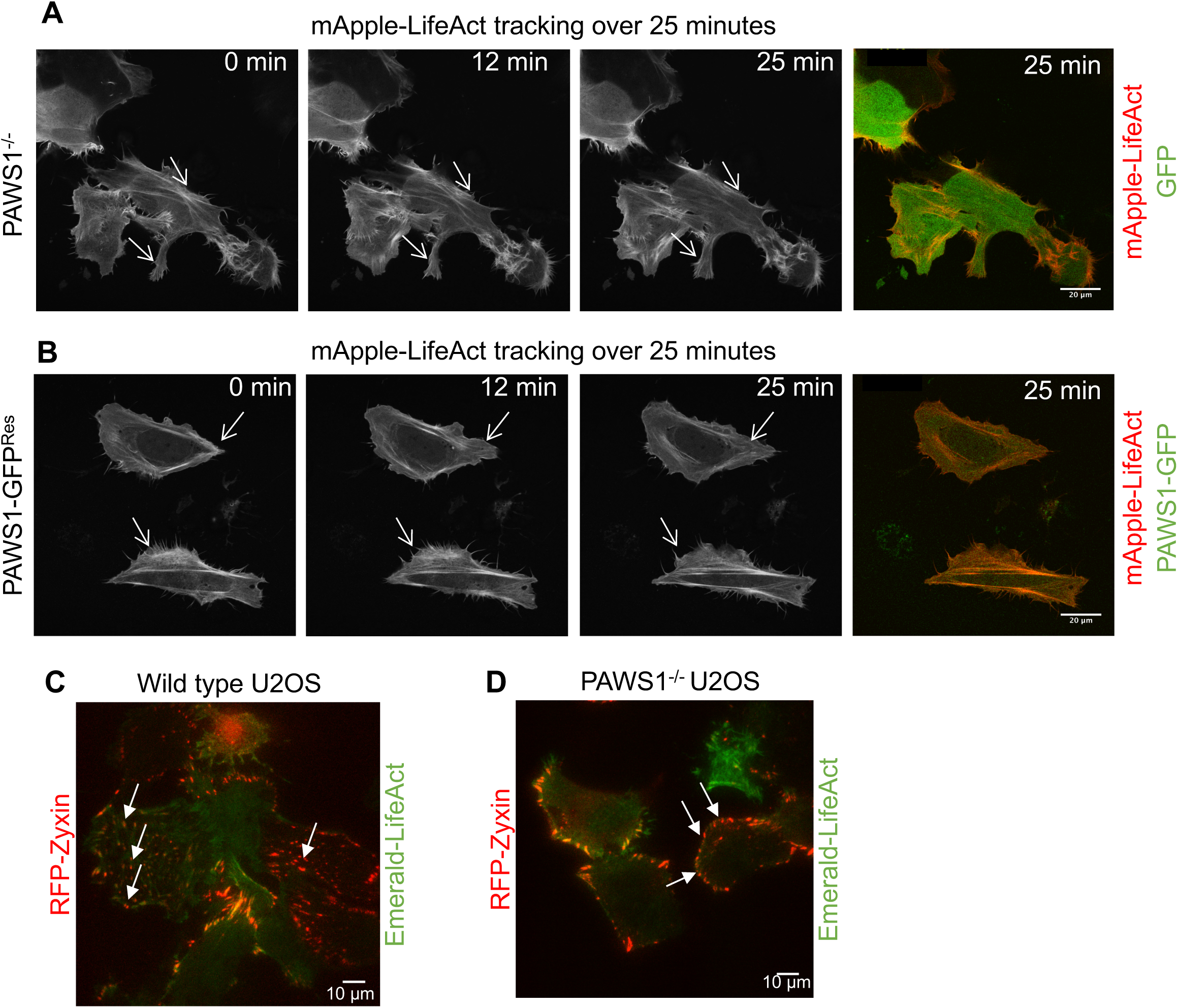
The effect of PAWS1 on actin and focal adhesion dynamics and distribution. **A.** GFP and LifeAct-mApple were transfected into PAWS1^-/-^ U2OS cells which were imaged for 25 minutes using a Zeiss confocal microscope at 60X magnification. Representative still images at the indicated times are presented. Scale bar is 20 µm. Static regions of membrane ruffles are indicated by the arrows. See Supplementary Movie 2 for actin dynamics over the time course of 25 minutes. **B.** As in A, except that PAWS1-GFP and mApple-LifeAct were transfected into PAWS1^-/-^ U2OS cells and imaged for 25 minutes. The dynamic ruffling of the membrane is indicated by the arrows. See Supplementary Movie 2 for actin dynamics over the time course of 25 minutes. **C.** U2OS wild type or **D.** PAWS1^-/-^ cells were transfected with RFP-Zyxin (punctate stains with arrows) and Emerald LifeAct, then imaged by TIRF microscopy at 60X magnifcation for 30 minutes to determine membrane dynamics of focal adhesions and cytoskeletal association. Scale bar is 10 µm.

Abnormal actin organization and cell shape can cause defects in cell migration(Schratt et al., 2002; Zaoui et al., 2008). To assess the role of PAWS1 in cell migration, we performed a lateral wound-healing assay(Huang et al., 2009). PAWS1^-/-^ and PAWS1^Res^ U2OS cells were cultured to confluency in adjacent chambers of a culture well divided by a small fixed-sized spacer, such that a uniform gap was created when the spacer was removed. Cell migration into the gap was monitored for up to 16h (Figure 1D). PAWS1^-/-^ cells migrated into the gap more slowly than PAWS1^Res^ cells at both 12 h and 16 h (Figure 1D). After 16 h, PAWS1^-/-^ cells showed 40% wound closure relative to the starting wound gap, compared with 85% for PAWS1^Res^ cells (Figure 1E). In a similar assay, live imaging of PAWS1^-/-^ and wild type U2OS cells on opposite sides of the wound showed that while wild type cells can form well-defined membrane ruffles and lamellipodia and migrate rapidly across the wound gap, PAWS1^-/-^ cells remain tightly connected to each other, form poorly-defined membrane ruffles and lamellipodia, and migrate slowly (Supplementary Movie 1). We also investigated the migration over time of wild type, PAWS1^-/-^ and PAWS1^Res^ cells towards a chemoattractant, after seeding cells in serum-free conditions on µ-Slide chemotaxis chambers with 10% FBS added on adjacent chambers as chemoattractant. Although no significant differences in the directionality of cell migration towards FBS were observed over the course of this assay, PAWS1^-/-^ cells displayed striking delay in adhesion compared to wild type or PAWS1^Res^ cells (Supplementary Figure 1). These observations indicate that PAWS1 plays a role in actin organization and cell migration in U2OS cells.

### Phenotypic characterization of PAWS1 actin defects

To understand how PAWS1 affects cytoskeletal dynamics, we first performed live cell imaging of PAWS^-/-^ U2OS cells transfected with either GFP control or PAWS1-GFP together with mApple-LifeAct (Figure 2A&B). PAWS1^-/-^ control cells displayed disorganized and static actin kinetics, suggesting that PAWS1 deletion causes defects in the organization and dynamics of the actin network (Figure 2A; Supplementary Movie 2A). In contrast, cells transfected with PAWS1-GFP had an organized and dynamic actin network, and membrane ruffling was observed throughout the 25 min imaging period (Figure 2B; Supplementary Movie 2B). Thus, the introduction of PAWS1-GFP in PAWS1^-/-^ cells was sufficient to restore membrane dynamics and the localization of actin in stress fibres (Figure 2B; Supplementary Movie 2B).

Focal adhesions anchor cell protrusions to the extracellular matrix. They organize the actin cytoskeleton and allow traction forces to be generated to move the cell body (Guo et al., 2006). We asked if focal adhesions are also affected by loss of PAWS1. Live-cell TIRF microscopy was carried out over 25 minutes on PAWS1^-/-^ U2OS or wild type cells transfected with the focal adhesion protein RFP-zyxin to determine focal adhesion dynamics and distribution (Fig 2C-D; Supplementary Movie 3). In wild type cells, zyxin displayed the expected punctate pattern throughout the basal surface of cells (Figure 2C). In contrast, PAWS1^-/-^ cells had a predominantly peripheral distribution of zyxin (Figure 2D), showing that focal adhesions fail to form properly. The defective distribution of zyxin in PAWS1^-/-^ cells was not observed in PAWS1^Res^ cells (Supplementary Figure 3).

**Figure 3.**
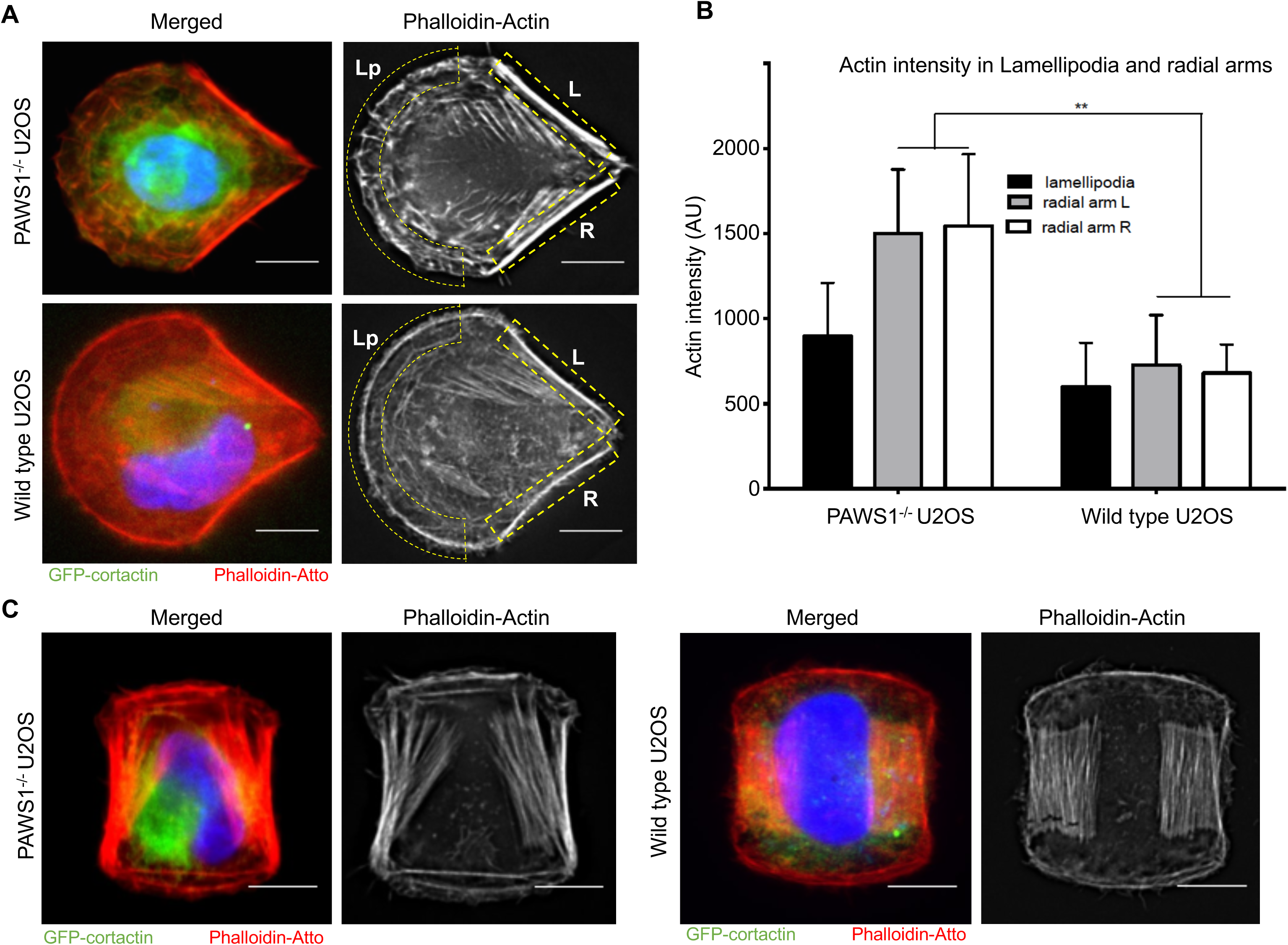
Micropattern cyclic strain analysis of PAWS1^-/-^ U2OS and wild type U2OS cells. **A.** PAWS1^-/-^ (upper panel) or wild type U2OS cells (lower panel) transfected with GFP-cortactin for 24 hours (as a secondary measure of membrane dynamics) and then stained with phalloidin-Atto and DAPI. Crossbow-micropattern chips were coated with 20 µg/ml fibronectin and cells were seeded at low density to allow adhesion of ~1 cell per pattern following gentle washing. Actin stress fibre organization was measured by widefield deconvolution microscopy. GFP-Cortactin (green), phalloidin-actin (red) and DAPI (blue). Scale bar is 20 µm **B.** Quantitation of 10 (WT) and 15 (PAWS1^-/-^) images of each cell genotype was performed using FIJI for accumulation of actin in the lamellipodia (Lp) and radial ventral arms at the trailing edge of the cells (indicated by L and R in yellow dashed boxes). Arbirtrary intensity units were measured and statistical Student’s t-test was conducted, allowing p<0.05 significance. **C.** As in A, except that double-crossbow H-patterned fibronectin-coated chips were used to plate PAWS1^-/-^ and wild type U2OS cells.

### Micropattern analysis of cytoskeletal actin fibres and cortactin in PAWS1^-/-^ U2OS cells

Bearing in mind the role of PAWS1 in cell morphology, migration, cytoskeletal organization and focal adhesion distribution, we decided to examine its contribution to the architecture of cortactin and actin fibres. To this end, PAWS1^-/-^ and control U2OS cells were plated onto fibronectin-coated crossbow and H-shaped (double-crossbow) micropatterns (Versaevel et al., 2016). We first noted that the lamellipodia of PAWS1^-/-^ cells plated on the ‘crossbow’ fibronectin micropattern had a disorganized actin pattern (Figure 3A; Supplementary Figure 4). Thus, in control cells there was a clearly defined continuous belt of actin that spanned the leading adhesive edge. However, in PAWS1^-/-^ cells we noted that this band was discontinuous and there were several spike-like actin projections (Figure 3A and Supplementary Figure 4). Control and PAWS1^-/-^ cells both showed the expected accumulation of actin along non-adhesive edges(Thery et al., 2006), but stress fibres between the adhesive regions of PAWS1^-/-^ cells were brighter than in control cells (Figure 3A&B). In the double crossbow micropattern, in addition to the defects observed above, we noted that stress fibres between the adhesion arms were not organised into proper parallel arrays in the PAWS1^-/-^ cells (Figure 3C and Supplementary Figure 5). There were no significant differences in the distribution of GFP-cortactin between WT and PAWS1^-/-^ cells in either micropattern, although the GFP-cortactin signal appeared to be more intense in PAWS1^-/-^ cells (Figure 3).

**Figure 4.**
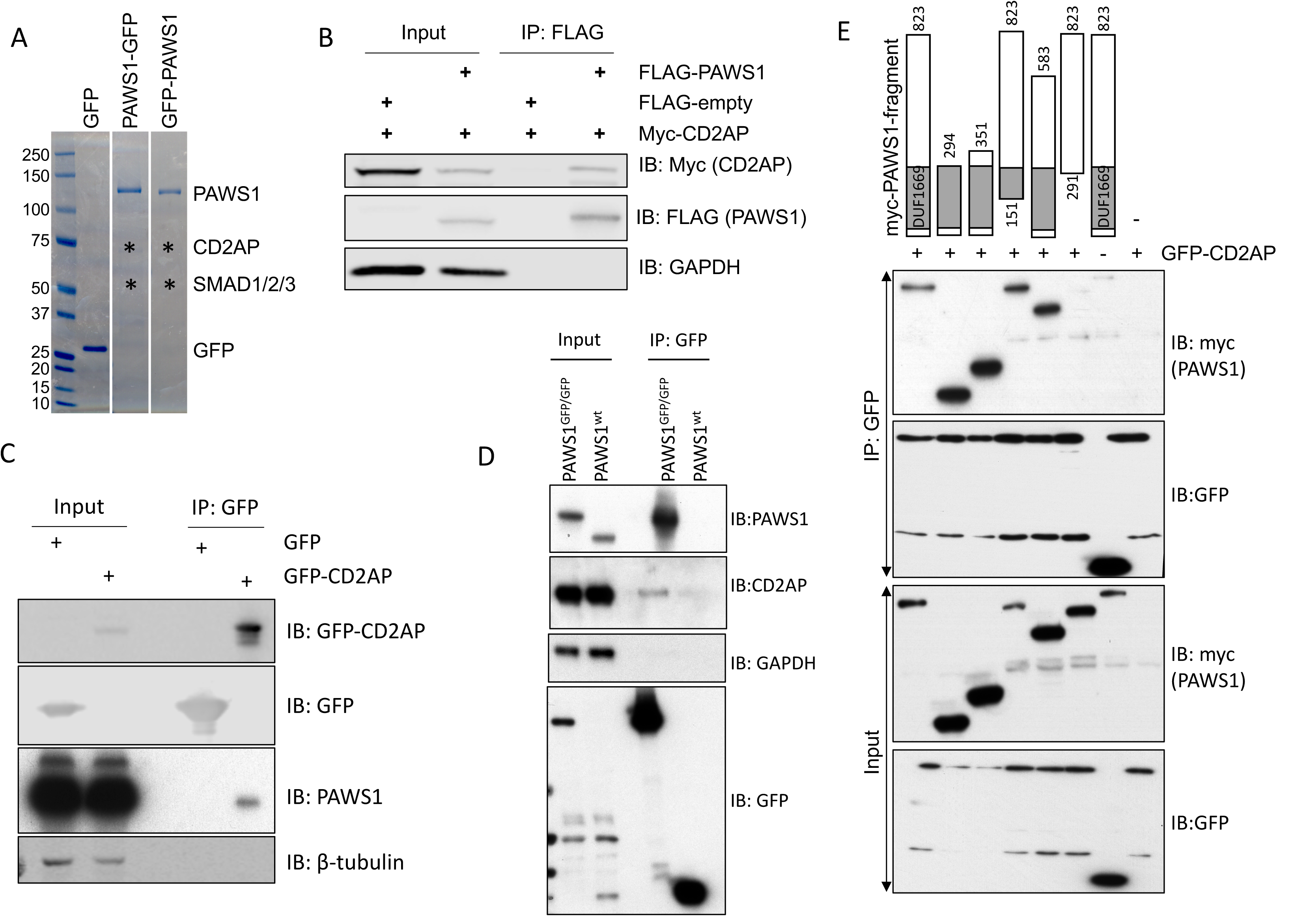
PAWS1 interacts with CD2AP. **A.** Anti-GFP IPs from HEK293 extracts expressing GFP alone, or PAWS1 tagged with GFP either at the C- or the N-terminus were resolved by SDS-PAGE and interacting proteins identified by mass spectrometry. The Coomassie stained gels indicating the approximate positions from where the designated interacting proteins were identified are included. **B.** Verification of interactions between Myc-tagged CD2AP and Flag-tagged PAWS1 by co-expression and IP experiments performed in PAWS1^-/-^ U2OS cells as indicated. **C.** Anti-GFP IPs from either GFP or GFP-CD2AP expressing cell extracts were subjected to immunoblotting with anti-GFP or anti-PAWS1 antibodies as indicated. **D.** Homozygous PAWS1-GFP knockin U2OS cells (PAWS1^GFP/GFP^) in which GFP-tag was introduced at the C-terminus of PAWS1 gene on both allelles using CRISPR/Cas9 and wild type U2OS cells transfected with GFP-control were subjected to anti-GFP-IPs. Extracts and anti-GFP IPs were then subjected to immunoblotting with anti-PAWS1 and anti-CD2AP as indicated. **E.** Mapping minimal PAWS1 region necessary for interaction with CD2AP. The indicated fragments of Myc-tagged PAWS1 were co-expressed with either GFP or GFP-CD2AP in PAWS1^-/-^ U2OS cells for 48 hrs. Extracts or anti-GFP IPs were subjected to immunoblotting with anti-myc and anti-GFP antibodies as indicated.

**Figure 5.**
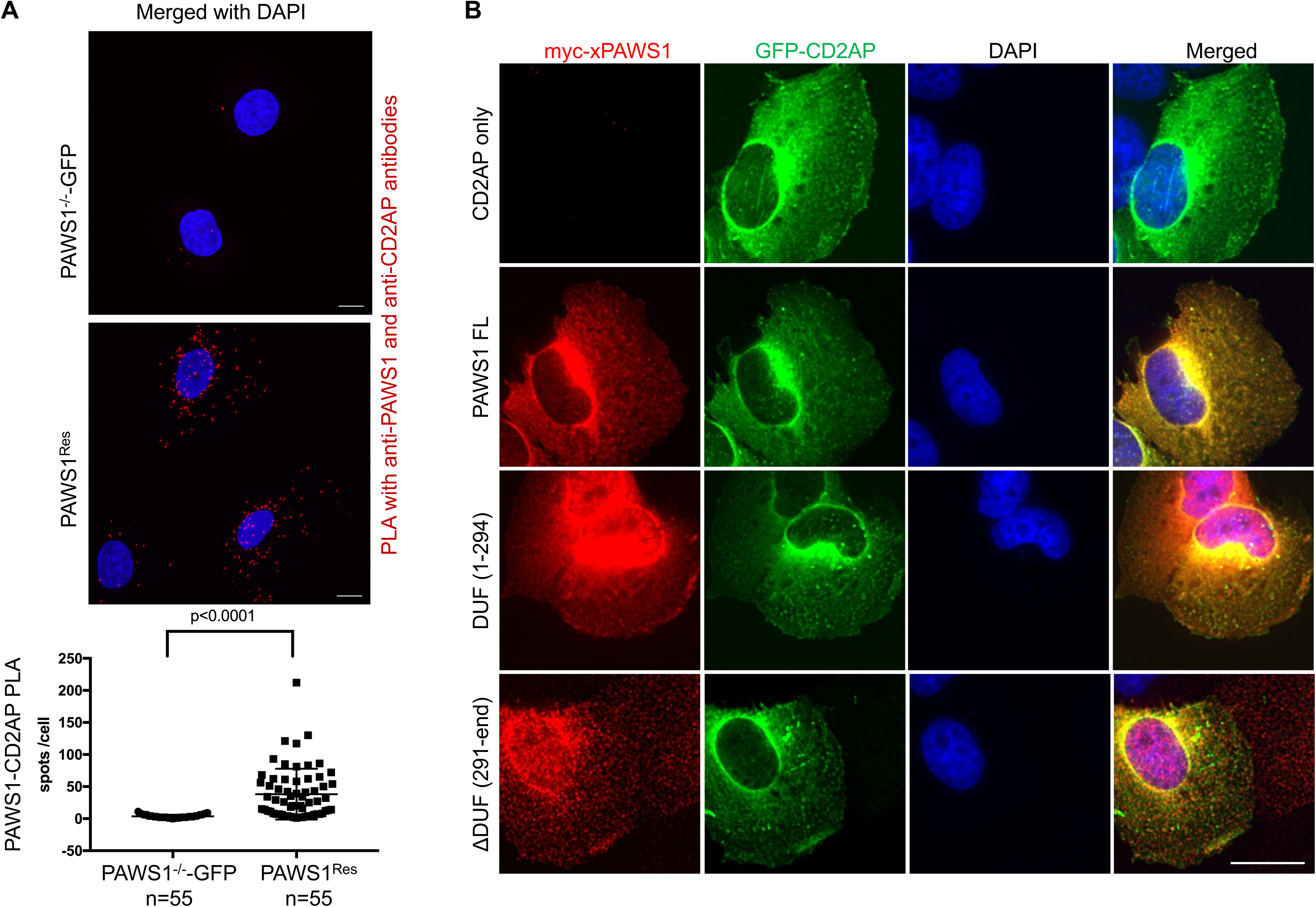
PAWS1 and CD2AP co-localize in U2OS cells. **A.** PAWS1^-/-^ U2OS cells restored with either control GFP (PAWS1^-/-^-GFP) or with WT PAWS1 (PAWS1^Res^) were plated onto 12-well Ibidi chamber slides. Cells were fixed and subjected to PLA using anti-PAWS1 and anti-CD2AP antibodies as described in the Methods section. Representative maximum intensity projections acquired using a Zeiss LSM710 confocal microscope using a Plan Apochromat 63x/1.4 oil lens captured using ZEN 2012 software are shown. PLA signals were counted using FIJI software and statistical analysis performed with Prism 7. Scale bar = 10 µm. **B.** PAWS1^-/-^ U2OS cells were transfected with GFP-CD2AP alone, or co-transfected with the indicated fragments of Myc-tagged xPAWS1. Cells were fixed in paraformaldehyde and subjected to immunofluorescence staining performed using anti-Myc-tag antibody, followed by Alexa Fluor (red) secondary antibody (ThermoFisher Scientific). Fluorescent images for Myc-PAWS1 (Red) and GFP-CD2AP (green) were captured using a DeltaVision system (Applied Precision). Z-series were collected at 0.2 μm intervals, and deconvolved using SoftWoRx (Applied Precision). Z-projections and image analysis were performed using OMERO (www.openmicroscopy.org). Scale bar = 20 μm.

### PAWS1 interacts with CD2AP, a key regulator of actin cytoskeleton

In order to understand the molecular mechanism by which PAWS1 modulates actin-cytoskeletal organization, we used mass spectrometry to identify PAWS1 interactors from tetracycline-inducible HEK293 cells(Yao et al., 2007; Yao et al., 1998) stably integrated with a single copy of either N-terminal or C-terminal GFP-tagged PAWS1 or GFP alone as control (Figure 4A). GFP-trap IPs of GFP alone, PAWS1-GFP and GFP-PAWS1 were resolved by SDS-PAGE and sections covering the entire lane for each sample were excised and digested with trypsin (Figure 4A). The resulting peptides were subjected to LC-MS/MS for identification. In addition to the SMAD isoforms, one of the most robust protein interactors identified for both GFP-PAWS1 and PAWS1-GFP but not GFP alone was CD2AP (Figure 4A). CD2AP plays a role in controlling actin cytoskeletal dynamics and cell migration(Srivatsan et al., 2013; Tang and Brieher, 2013). We went on to verify the interaction between PAWS1 and CD2AP. Upon co-expression in HEK293 cells, Myc-CD2AP is detected in FLAG-PAWS1 immunoprecipitations but not in control FLAG IPs (Figure 4B). Endogenous PAWS1 was detected in GFP-CD2AP IPs but not in control GFP IPs from U2OS cells transiently transfected with either GFP-CD2AP or GFP (Figure 4C). In order to verify endogenous interaction between PAWS1 and CD2AP, in the absence of robust PAWS1 and CD2AP immunoprecipitating antibodies, we generated homozygous PAWS1-GFP knockin U2OS cells using CRISPR/Cas9 (Figure 4D). Endogenous CD2AP and PAWS1 were detected in anti-GFP IPs from these cells but not wild type cells transfected with GFP control (Figure 4D). Together, these observations suggest that there is a robust interaction between PAWS1 and CD2AP. To map the PAWS1 interaction domain, we co-expressed Myc-tagged PAWS1 fragments with full length GFP-CD2AP in PAWS1^-/-^ cells and performed co-IP experiments (Figure 4E). GFP-CD2AP co-precipitated PAWS1 only if PAWS1 contained residues 151-291 within the *DUF1669* domain (Figure 4E). Consistent with these observations, when ~100 amino acid fragments of FLAG-tagged PAWS1 spanning the entire protein were co-expressed with full-length Myc-tagged CD2AP in PAWS1^-/-^ cells, only FLAG-PAWS1(204-294) and full-length FLAG-PAWS1 were able to co-immunoprecipitate Myc-CD2AP (Supplementary Figure 6).

**Figure 6.**
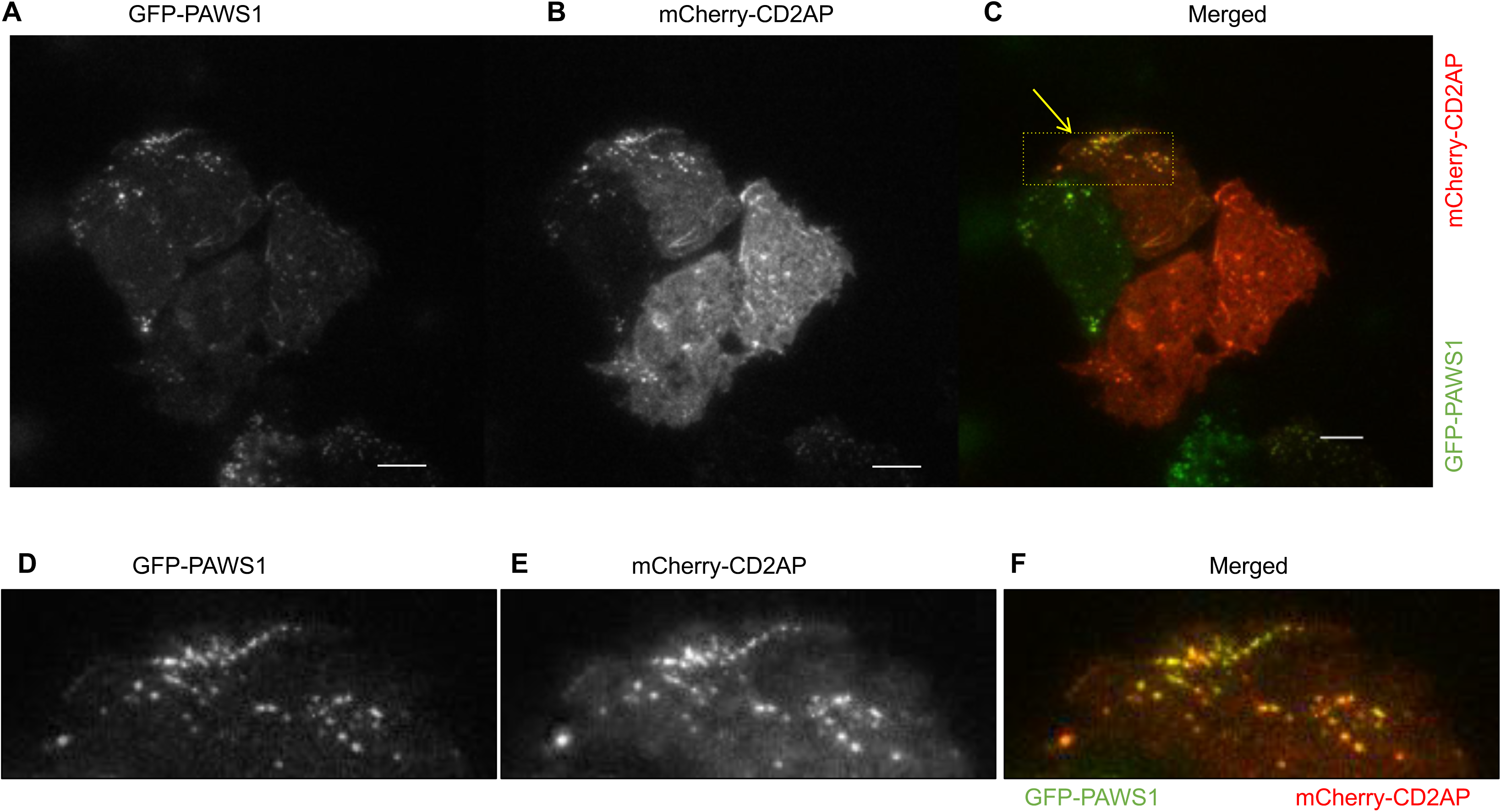
CD2AP and PAWS1 co-localize in dynamic punctate structures proximal to the plasma membrane in wild type U2OS cells. Total internal reflection fluorescence microscopy (TIRF) in U2OS cells transfected with mCherry-CD2AP and GFP-PAWS1.A. GFP-PAWS1 **B.** mCherry-CD2AP and **C.** Merged mCherry-CD2AP and GFP-PAWS1 indicate co-localization around the plasma membrane. Scale bar is 10 µm. **D-F.** Equivalent images from the zoomed in boxed-region indicated by an arrow in **C.**

### PAWS1 interacts and co-localizes with CD2AP in cells

To further confirm the interaction between CD2AP and PAWS1 in cells, we performed an *in situ* proximity ligation assay (PLA) (Soderberg et al., 2006). PLA technology uses species specific secondary antibody probes that are labelled with unique DNA tags. If the probes bind to their respective targets in close proximity to one another (within 40 nm) their DNA tags can be ligated and used as a template for a rolling circle amplification reaction. When supplemented with a fluorescent label, the reaction produces fluorescent puncta, which is evidence of specific protein-protein interactions. In control PAWS^-/-^-GFP cells only a few scattered PLA puncta per cell were observed (Figure 5A). In contrast, in the PAWS1^Res^ cells we observed a significant number of CD2AP-PAWS1 specific puncta (Figure 5A) demonstrating that CD2AP and PAWS1 interact *in situ*. Similarly, through immunostaining and fluorescence microscopy on fixed cells, we assessed the subcellular localization of GFP-CD2AP and myc-PAWS1 fragments co-expressed in PAWS1^-/-^ U2OS cells (Figure 5B). In the absence of PAWS1, GFP-CD2AP was localized predominantly in the cytoplasm. When co-expressed, a substantial overlapping cytoplasmic staining was observed for both full length PAWS1 and GFP-CD2AP (Figure 5B). The DUF1669 domain (1-294) of PAWS1, which binds CD2AP (Figure 4E), showed both nuclear and cytoplasmic staining but the overlapping staining with CD2AP was only observed in the cytoplasm (Figure 5B), suggesting CD2AP co-localizes with PAWS1 only in the cytoplasm. When GFP-CD2AP was co-expressed with an interaction deficient PAWS1(291-end) fragment (Figure 4E), very little overlapping staining was observed (Figure 5B). A very distinct pan-cellular punctate staining for PAWS1(291-end) fragment was observed (Figure 5B). Similar distributions of PAWS1-CD2AP interactions were observed in the *in situ* PLA assays (Figure 5). Together with the immunoprecipitation experiments, these data suggest robust interactions between CD2AP and PAWS1.

Next, in order to investigate the dynamics of PAWS1-CD2AP interaction in cells, we used live cell TIRF microscopy on wild type U2OS cells transfected with GFP-PAWS1 and mCherry-CD2AP. In addition to cytoplasmic co-localization, under these conditions we observed that the two proteins co-localize in dynamic punctate structures adjacent to ruffling membranes and lamellipodia (Figure 6; C&F for merged; Supplementary Movie 4). Over the course of live cell imaging, some non-overlapping, predominantly cytoplasmic staining of both GFP-PAWS1 and mCherry-CD2AP was also observed (Figure 6). The dynamic co-localization of PAWS1 and CD2AP in distinct structures suggests there might be regulated interaction between these proteins. Interestingly, when the co-localization of transiently transfected GFP-PAWS1 and mCherry-CD2AP was explored using TIRF microscopy in PAWS1^-/-^ U2OS cells, similar overlapping punctate structures adjacent to ruffling membranes were observed (Figure 7A-C), but in an adjacent PAWS1^-/-^ cell in which GFP-PAWS1 was absent, no punctate structures were visible for mCherry-CD2AP (Figure 7A-C). These observations suggest that PAWS1 may be required for localization of CD2AP at the dynamic punctate structures.

**Figure 7.**
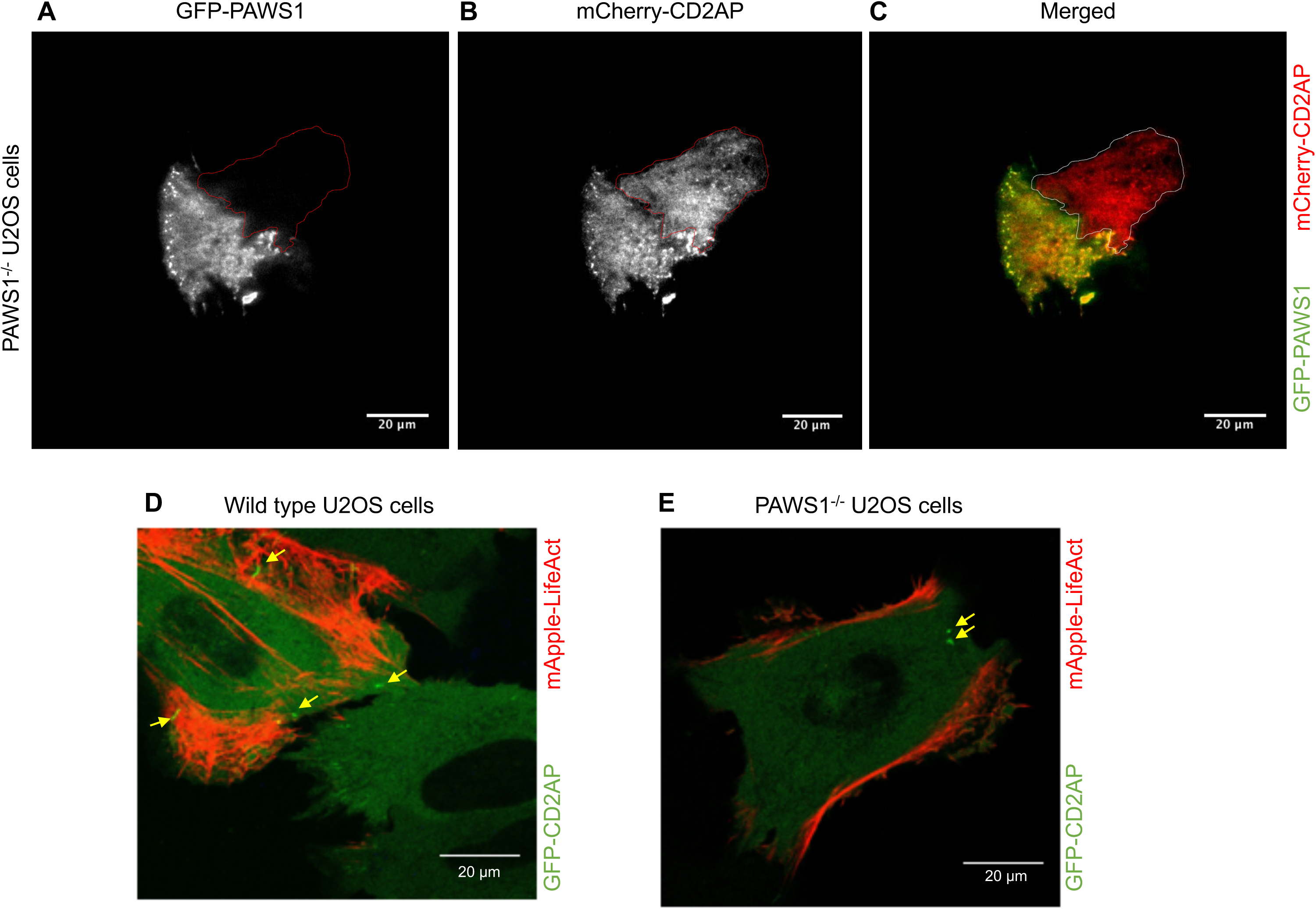
PAWS1 appears to regulate localization of CD2AP with dynamic actin. **A-C.** PAWS1^-/-^ U2OS cells transfected with mCherry-CD2AP (red) and GFP-PAWS1 (green) for 24 hours were followed by TIRF live cell imaging for 25 minutes to assess the localization and dynamics of CD2AP and PAWS1 around the membranes. Note that cross-channel bleed-through is minimal, indicated by lack of cross channel GFP-excitation/emission in only mCherry-positive cell outlined. Representative images are shown. Scale bars indicate 20 µm. **D.** Wild type U2OS and **E.** PAWS1^-/-^ U2OS cells were transfected with GFP-CD2AP (green) and mApple-LifeAct (red) for 24 hours followed by wide-field fluorescence microscopy live cell imaging at 40X magnification for 30 mins. Representative images are shown. GFP-CD2AP puncta are indicated by white arrows. Scale bars indicate 20 µm.

To understand the impact of PAWS1 on CD2AP localization in the context of actin cytoskeletal dynamics, we analysed wild type and PAWS1^-/-^ U2OS cells transfected with GFP-CD2AP and mApple-LifeAct by wide-field fluorescence microscopy (Figure 7D&E). In wild type U2OS cells, GFP-CD2AP puncta were visualized close to the active ruffling lamellipodia and actin-rich components of the plasma membranes (Figure 7D). In contrast, in PAWS1^-/-^ cells GFP-CD2AP was distributed diffusely in the cytoplasm and any visible puncta did not extend into lamellipodia (Figure 7E).

## Discussion

Our previous work has shown that PAWS1 interacts with SMAD1, that it is a substrate of type I BMP receptor kinases, and that it is involved in Smad4-independent BMP signalling. We also demonstrated that PAWS1 regulates the expression of several non-BMP target genes, suggesting that it has roles beyond the BMP pathway. By knocking out PAWS1 from U2OS osteosarcoma cells, we show here that PAWS1 plays a role in actin organization, morphology and migration in U2OS cells, and that it is likely to exert these effects through its interaction with CD2AP. In particular, the interaction of PAWS1 with CD2AP at the cell periphery appears to control actin dynamics to initiate lammellipodia formation and cellular migration. Future work will explore the ability of PAWS1 to influence both BMP signalling and the actin cytoskeleton, to ask whether the two functions are linked and to ask whether any other activities can be attributed to this protein.

## Materials and Methods

### Antibodies

An anti-FAM83G antibody was produced by the Dundee Division of Signal Transduction and Therapy (DSTT) as a sheep polyclonal antibody against the C-terminus of FAM83G (S876C); sheep anti-GFP was also produced by the DSTT (S268B). Other antibodies used in these studies were: Myc-tag (CST); FLAG-M2-HRP (Sigma); MYC-HRP (Roche); CD2AP (Gift from A. Shaw), CD2AP (clone 2A2.1 Millipore), anti-FAM83G (HPA023940, Sigma), and actin (Abcam). Secondary Rabbit, Mouse and Sheep antibodies conjugated to horseradish peroxidase were used at 1:10,000 (Santa Cruz Biotech). Fluorophore conjugated phalloidin (Alexa 488 and Atto 562) and Alexa-Fluor-594 anti-mouse (ThermoFisher) were used for fluorescence microscopy.

### Cell culture

Cells (U2OS, 293T, and HEK293) were cultured in Dulbecco’s modified Eagle’s medium (DMEM; Gibco) with 10% fetal bovine serum (Hyclone), 1% penicillin/streptomycin (Lonza) and 2 mM L-Glutamine (Lonza). 293T cultures were supplemented with sodium pyruvate after onset of retrovirus production. All cells in culture were routinely tested for mycoplasma contamination and verified as mycoplasma-negative.

### Vectors

Plasmids were designed and cloned in the DSTT and site-directed mutagenesis was used to generate mutant forms of PAWS1. Other plasmids used were pBABE-PAWS1-puro, pBABE, pCDNA5-frt-TO-nGFP-PAWS1, pCDNA5-frt-TO −PAWS1-cGFP, pCDNA5-frt-TO-GFP, pCMV-GFP-CD2AP, mCherry-xCD2AP, GFP-CD2AP, mApple-LifeAct (Life Technologies), Emerald-LifeAct (Life Technologies), RFP-zyxin, pCS2-xPAWS1, pCS2-xCD2AP, pCDNA-PAWS1-FLAG. All DNA constructs were verified by DNA sequencing, performed by the DNA Sequencing & Services (MRCPPU, College of Life Sciences, University of Dundee, Scotland, http://www.dnaseq.co.uk) using Applied Biosystems Big-Dye Ver 3.1 chemistry on an Applied Biosystems model 3730 automated capillary DNA sequencer. All constructs are available to request from the MRC-PPU reagents webpage (http://mrcppureagents.dundee.ac.uk).

### FAM83G knockout using CRISPR/Cas9 genome editing

CRISPR/Cas9 mediated deletion of FAM83G/PAWS1 in osteosarcoma cells (U2OS) was performed using Cas9 and a single gRNA targeting approach to delete exon 2 of the RefSeq gene for FAM83G (NM_001039999.2). Vectors containing the Cas9 and FAM83G targeting gRNA (ggaccgctccatcccgcagctgg) were transfected into 1x10^6^ U2OS cells followed by selection with 2 ug/ml puromycin and single cell sorting to isolate clone candidates with gene deletion. Sequencing of the gRNA targeting region indicated a 5-bp deletion causing a frameshift in FAM83G gene.

### Retroviral FAM83G/PAWS1 expression

Retroviral constructs of pBABE-puromycin, pBABE-PAWS1, or pBABE-GFP (5 µg each) were co-transfected with pCMV-gag/pol (4.5 µg) and pCMV-VSVG (0.5 µg) using Polyethylenimine (PEI, 1mg/ml; 25 µl) in 1 ml OPTIMEM low serum medium into a 10-cm dish of HEK293T. After 40 hours of culture, supernatant medium was filtered (0.45 µm) and applied to recipient cells and supplemented with 8 µg/ml polybrene (Sigma #H9268, Hexadimethrine bromide). Recipient U2OS cells were plated at 40-50% confluence and then infected with the indicated virus for 24 hr. Following virus infection, U2OS cells were treated with puromycin at 2 µg/ml to select for vector integration by the virus.

### 2-Dimensional Lateral cell migration

U2OS cells were plated into iBIDI insert chambers (Cat# 80209) 18 hours before 2-dimensional migration assays were performed. Equal numbers of 40-60,000 cells were plated on both sides of the chamber and the silicone insert was removed to allow lateral migration. Cells were incubated in a 5% CO_2_ regulated and 37°C temperature controlled chamber. Images were collected for 18-24 hrs using a Nikon Eclipse Ti microscope. Images of the wound gap were collected every 5 minutes by a Photometrics Cascade II CCD camera with Nikon NIS elements software. Wound closure was measured by Image J and reported as a percentage of closure relative to the starting wound size.

### Chemotaxis Assay

U2OS^WT^, PAWS1^-/-^ and PAWS1^Res^ cells were trypsinized and introduced into one end of a µ-Slide chemotaxis chamber (iBidi, Cat#80326) at 3x10^6^ cells/ml, while the opposite end was loaded with medium containing 10% FBS (Pepperell and Watt, 2013). Images of migrating cells were collected every 5 minutes using a Nikon Eclipse Ti microscope and Photometrics II CCD camera with Nikon NIS elements software as above.

### F-actin staining

U2OS cells were seeded onto microscope slides at low density and allowed to grow to 20-30% confluence. Cells were then fixed with 4% paraformadlehyde (PFA) for 30 minutes, and washed 2X in DMEM/HEPES pH 7.4 followed by 10 min incubation in DMEM/HEPES. Cells were washed 1X in PBS then 0.2% Triton X100 in PBS for 3-5 min. Cells were washed in 1% BSA/PBS followed by staining with Phalloidin (Alexa-Fluor-488 or Atto-562) performed at 1:500 dilution in the dark for 1 hour at room temperature. Following incubation, slides were washed 3X in BSA/PBS solution. Coverslips were mounted in Prolong gold with DAPI for nuclear staining. Coverslips were allowed to dry briefly then sealed and imaged by widefield deconvolution microscopy.

### Immunofluorescence

Cells were fixed with 4% PFA in PBS for 10 min, and permeabilised with 0.5% Triton X-100 in PBS for 5 min. Coverslips were incubated in blocking buffer (3% bovine serum albumin, 0.1% Triton X-100 in PBS) for 30 min, followed by primary antibodies diluted in blocking buffer for 1 h. Cells were washed with 0.1% Triton X-100 in PBS, and incubated with the appropriate Alexa Fluor secondary antibodies (ThermoFisher Scientific). Images were captured using a DeltaVision system (Applied Precision). Z-series were collected at 0.2 μm intervals, and deconvolved using SoftWoRx (Applied Precision). Z-projections and image analysis were performed using OMERO (www.openmicroscopy.org).

### Proximity Ligation Assay (PLA)

PLA (DuoLink, Sigma) was performed following the manufactures’ recommendations. In brief, cells plated in 12 well Ibidi chamber slides (cat #81201) were fixed in 4% formalin in PBS for 10 min and then permeabilised in 0.5% TX100 for 5 min. Cells were blocked in PLA blocking solution for 30 min at 37˚C followed by incubation with primary antibodies diluted in antibody dilution solution (1:500 mouse anti-CD2AP (Millipore), 1:2000 rabbit anti-FAM83G (Sigma)) for 1 hr at RT. Cells were then washed twice for 5 min in wash buffer A and then incubated for 1 hr at 37 ˚C with PLA probes (rabbit (+) and mouse (–) diluted in antibody dilution solution). Following two washes in buffer A, the probes were ligated at 37 ˚C for 30 min and then fluorescently labelled with orange reagent during a 100 min amplification reaction at 37 ˚C. Cells were then washed twice for 10 min in buffer B followed by a brief 1 min wash in 0.01x buffer B. Cells were mounted using Duolink In Situ Mounting Medium with DAPI and the coverslip sealed with nail polish. Images were acquired with a Zeiss LSM710 confocal microscope using a Plan Apochromat 63x/1.4 oil lens. Z-stacks (6.56 µm) were captured using ZEN 2012 software. PLA signals were counted using FIJI software and statistical analysis performed with Prism 7.

### Transfection of fluorescent proteins

Cells were transfected with 2-5 µg of GFP-PAWS1, mCherry-CD2AP, RFP-Zyxin (a gift from Yu-li Wang), mApple-LifeAct (Invitrogen) or Emerald LifeAct (Invitrogen) along with Fugene HD or PEI. Cells were cultured for 24-48 hours and then imaged or processed as indicated.

### Live cell imaging

U2OS cells were plated onto polystyrene CellView Culture (Greiner Bio-One) glass bottom dishes. Following transfection, images were captured using a Zeiss LSM 700 confocal microscope in a regulated chamber with 5% CO2 at 37°C for 1-2 hours as indicated. Images were taken of each fluorophore in sequence at 5 minute increments using Zen software.

### Total Internal Reflection Fluorescence microscopy (TIRF): Live cell imaging

TIRF microscopy was used to detect interactions between FAM83G/PAWS1 as well as CD2AP and actin cytoskeleton at the plasma membrane. Cells were plated in World Precision Instrument imaging chambers and transiently transfected with fluorophore tagged PAWS1, CD2AP, LifeAct actin trackers (mApple or mEmerald) and RFP-zyxin and imaged in CO2-independent medium (Leibovitz’s L-15; Life Technologies). TIRF was performed on a Nikon Ti-U microscope with an environmental control chamber (Okolab, Pozzuoli, Italy), a PAU/TIRF slider, 63x and 100x 1.49 N.A. objectives, PerfectFocus system, a custom-built four-color (405nm, 488nm, 561nm, 647nm) diode laser (Coherent Inc., Santa Clara, CA, USA) system with a Gooch and Housego (Ilminster, UK) AOTF shutter (Solamere Technology; Salt Lake City, UT, USA), an emission filter wheel (Nikon) with appropriate filters for eliminating crosstalk between channels (Chroma Technology Corp, Bellow Falls, VT, USA) and a Photometrics Evolve Delta camera (Tucson, AZ, USA). Images were all captured with µ-Manager (Open Imaging Inc., San Francisco, CA, USA).

### Micropattern Analysis and Wide-field Fluorescence Microscopy

Micropattern chips were from CYTOO (Grenoble, France) in multi-shape patterns including crossbow, H-pattern and Y-pattern. CYTOO 22-chip was coated with 20 µg/ml fibronectin in PBS for 2 hours at room temperature according to the manufacturer’s recommendation. The chip was washed 3 times in PBS and then air dried overnight at 4°C. U2OS cells were split and plated onto the chip then washed after 1 hour of attachment to minimize cytophobic surface binding. Cells were then fixed, stained with phalloidin-Alexa 562/594 and DAPI, and Z-stacks were collected on a widefield deconvolution microscope (GE Healthcare Life Sciences) or LSM 700 confocal microscope. Image analysis was performed with Image J on images acquired with equivalent exposure times for each experiment. F-actin accumulation in 10-15 cells from PAWS1^+/+^ or PAWS1^-/-^ U2OS cells was quantified in the lamellipodia and the radial arms in the ‘trailing ventral arms’ of the cells fixed on crossbow micropatterns. Cells also attached and formed stress fibers in the H pattern, and images were collected and analyzed in a similar manner.

### Cell Lysis, Affinity Purification and Western Blotting

Cells were washed twice in ice-cold PBS then scraped on ice into lysis buffer (50 mM Tris–HCl pH 7.5, 0.27 M sucrose, 150 mM NaCl, 1 mM EGTA, 1 mM EDTA, 1 mM sodium orthovanadate, 1 mM sodium β - glycerophosphate, 50 mM sodium fluoride, 5 mM sodium pyrophosphate, 1% (v/v) Triton X-100 and 0.5% Nonidet P-40) supplemented with complete protease inhibitor cocktail tablet (Roche). Lysates were clarified by centrifugation at ~17,000x g at 4°C. Protein concentration was estimated using a Bradford assay (ThermoFisher). Typically, 15-30 µg was used for SDS-PAGE and 250 µg-1 mg extract proteins was used for immunoprecipitation and interaction studies. For immunoprecipitation, extracts were loaded with 10 µl of GFP trap beads (ChromoTek) or anti-FLAG-M2 beads (Sigma) and incubated on a rotator for 4-16 hours at 4°C. Beads were washed in lysis buffer including 0.25M NaCl once, followed by lysis buffer. Purified proteins were eluted in 1X sample buffer (50mM Tris–HCl pH 6.8, 10% SDS, 50% glycerol, 0.1% bromophenol blue; 0.1% β-mercaptoethanol) and heated to 95°C for 5 minutes, and 25-50% of sample was fractionated using 4-20% or 10% SDS-PAGE as indicated. Gels were electroblotted to Immobilon PVDF (Millipore) and blocked in 5% milk with TTBS (50 mM Tris–HCl pH 7.5, 0.2% Tween-20, 150 mM NaCl,) for 1 hour at room temperature. Immunoblotting was performed with antibody at 1 µg/ml overnight in either 5% milk-TTBS or 5% BSA-TTBS at 4°C on a shaker. Blots were washed 4X in TTBS and probed with secondary antibody to rabbit, sheep or mouse conjugated with HRP (horseradish peroxidase) (Santa Cruz Biotech) at 1:10,000 dilution in 5% milk-TTBS for 1 hour at room temperature. Membranes were then washed 4X with TTBS followed by enhanced chemiluminescent detection (ThermoFisher) and exposure to X-ray film or on a Gel Doc XR+ system using Image Lab software.

### Mass spectrometry Analysis

GFP, GFP-PAWS1 or PAWS1-GFP constructs integrated stably into 293 TRex cells (Invitrogen) were expressed upon treatment with 20 ng/ml doxycyline for 16 h. Proteins were affinity purified by GFP-trap beads (ChromoTek) and subjected to mass-spectrometry analysis was performed as previously(Vogt et al., 2014). Briefly, purified proteins were separated by 4-12% gradient SDS-PAGE then stained with colloidal Coomassie blue overnight. The gel was washed in distilled H2O until background staining was minimal. Six gel pieces covering the entire lanes for each pulldown were excised, trypsin digested, and peptides prepared for HPLC gradient fractionation and elution into a Thermo Scientific Velos Orbitrap mass spectrometer. Ion assignments were conducted by in-silico Mascot scoring (www.matrixscience.com) and peptide protein assignments were reported in Scaffold 4.1 (www.proteomesoftware.com).

## Acknowledgements

We thank Dr. A. Shaw (Genentech) for anti-CD2AP antibody. We thank L. Fin, J. Stark, A. Muir and K. Airey for help with tissue culture, the staff at the Sequencing Service (School of Life Sciences, University of Dundee, UK) for DNA sequencing, and the protein & antibody production and cloning teams at the Division of Signal Transduction Therapy (DSTT; University of Dundee) coordinated by H. McLauchlan and J. Hastie.

## Author contributions

TDC performed most of the experiments, collected and analysed data and wrote the initial draft manuscript and edited subsequent drafts. KW, PB, KSD performed some experiments and analysed data. TM designed strategies and developed methodologies for and generated CRIPSR/Cas9 knockout and knock-in constructs. NW performed cloning and mutagenesis for most of the constructs. AP and EG assisted with microscopy design, set-up, imaging and data analysis. JCS contributed with data analysis and the composition of the manuscript. GPS conceived the project and wrote the manuscript.

---

**Supplementary Figure 1.**
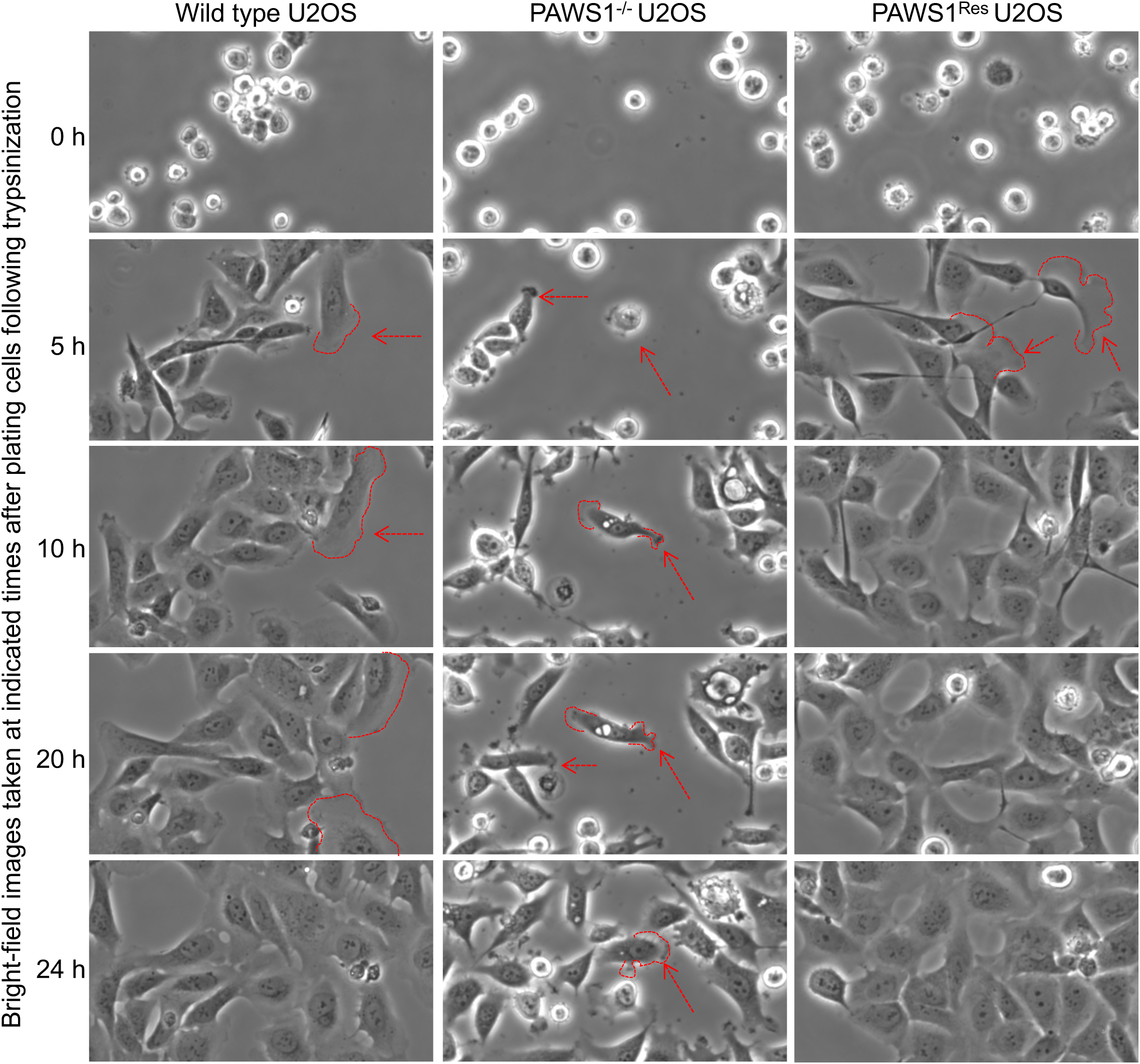
Chemotaxis assay for wild type U2OS, PAWS1^-/-^ U2OS and PAWS1^Res^ U2OS cells with FBS. Freshly trypsinized cells were introduced into a µ-Slide chemotaxis chamber at one end at 3x10^6^ cells/ml while the opposite end (right side of the images) was loaded with DMEM medium containing 10% FBS. Images were acquired every 5 hours by a Photometrics II CCD camera with Nikon NIS elements software. Outlined in red are cells developing lamellipodia or pseudo-filopodia. PAWS1^-/-^ U2OS cells do not fully develop lamellipodia.

**Supplementary Figure 2.**
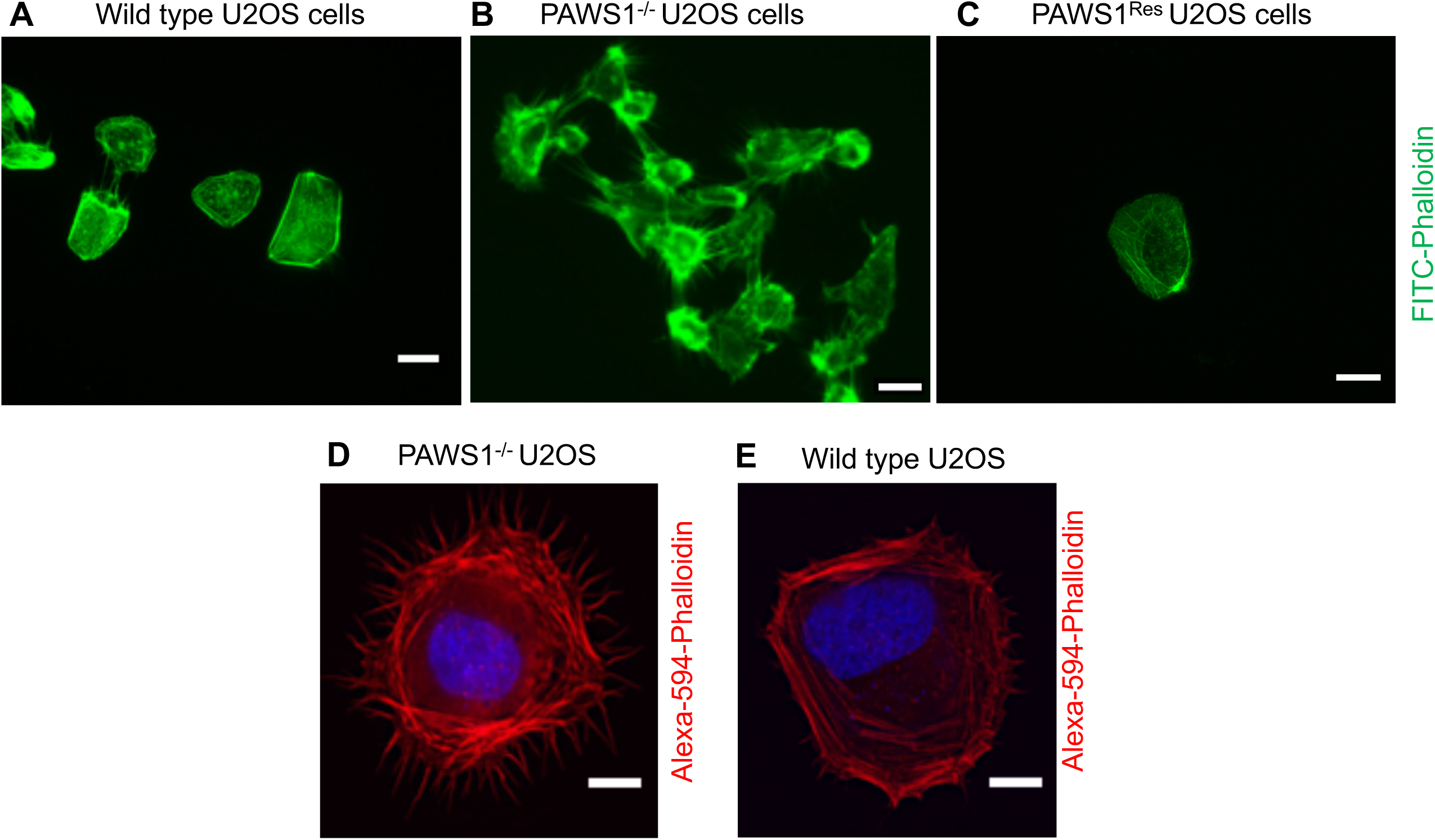
Analysis of actin fibres in wild type and PAWS1 ^-/-^ U2OS cells. **A.** wild type U2OS **B.** PAWS1^-/-^ U2OS and **C.** PAWS1^Res^ U2OS cells were fixed and stained with FITC-Phalloidin (green). Representative images are included. Scale bars indicate 10 µm. **D.** PAWS1^-/-^ U2OS cells and **E.** wild type U2OS were fixed and stained with Alexa-594-Phalloidin (red). Representative images are included. Scale bars indicate 10 µm.

**Supplementary Figure 3.**
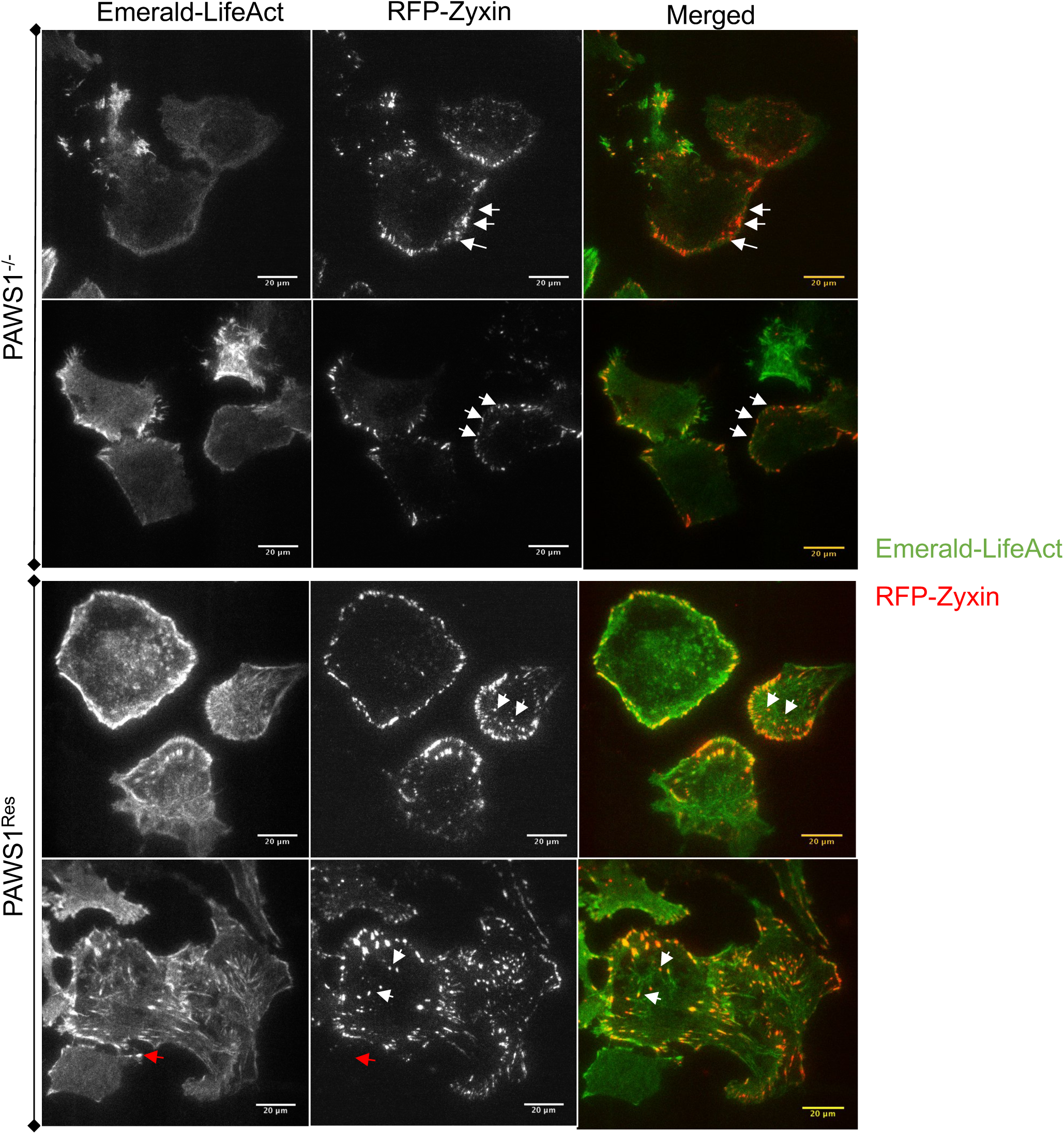
Analysis of focal adhesion dynamics in PAWS1^-/-^ and PAWS1^Res^ cells using TIRF microscopy. PAWS1^-/-^(upper panel) andPAWS1^Res^(lower panel) U2OS cells were transfected with RFP-zyxin (red) and Emerald-LifeAct (green) to measure changes in cytoskeletal and focal adhesion dynamics. Arrows indicate focal adhesion points. Panels are duplicate representative images. TIRF microscopy was conducted for the analysis of membrane dynamics of the adhesion process.

**Supplementary Figure 4.**
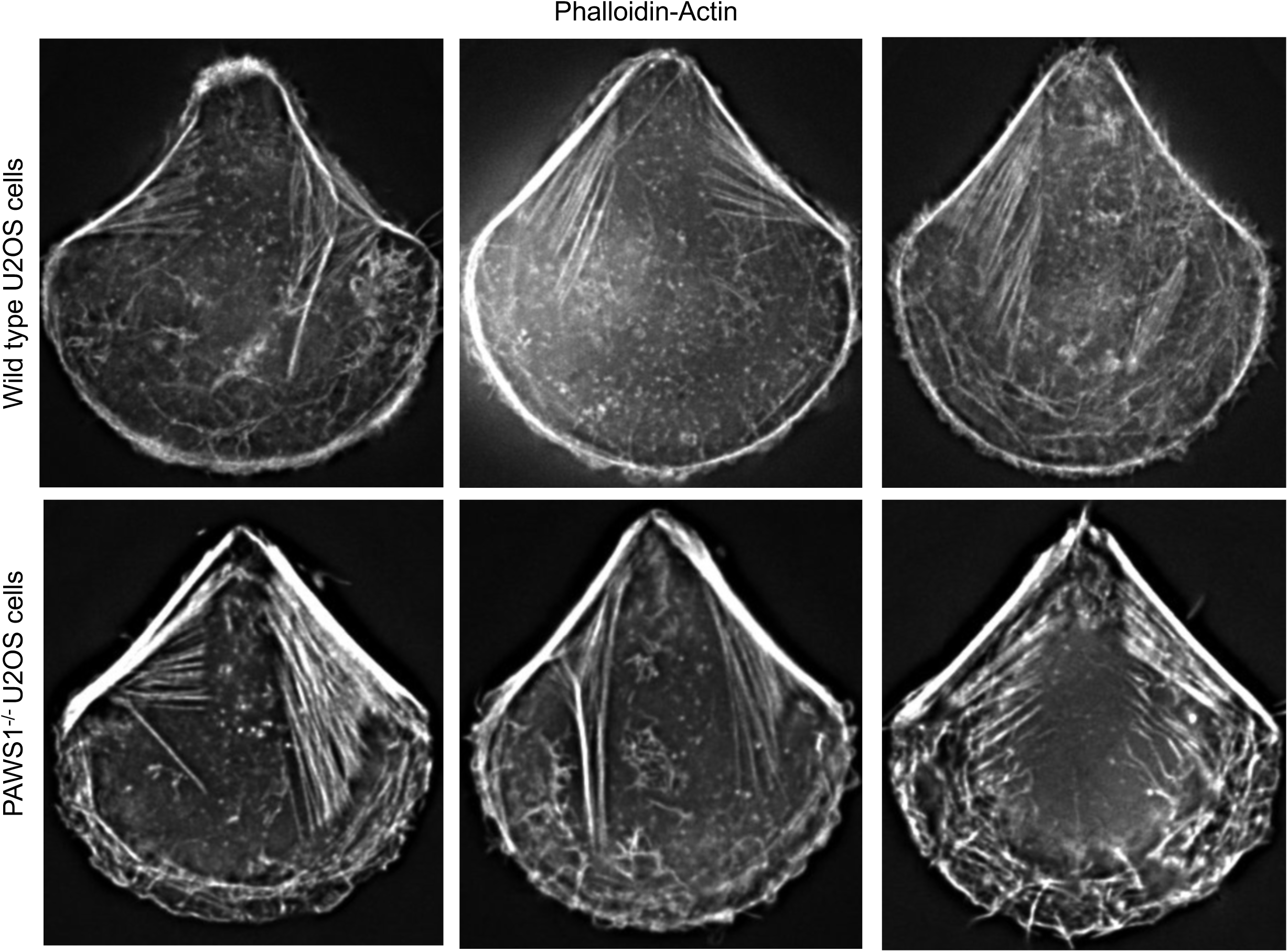
Additional representative images of micropattern adhesion dynamics of phalloidin-stain in wild type and PAWS1^-/-^ U2OS cells in the crossbow pattern. Monochrome images of each genotype are presented to indicate the shape and structure of actin fibres. Cells and images were processed as described in Legends to Figure 3A.

**Supplementary Figure 5.**
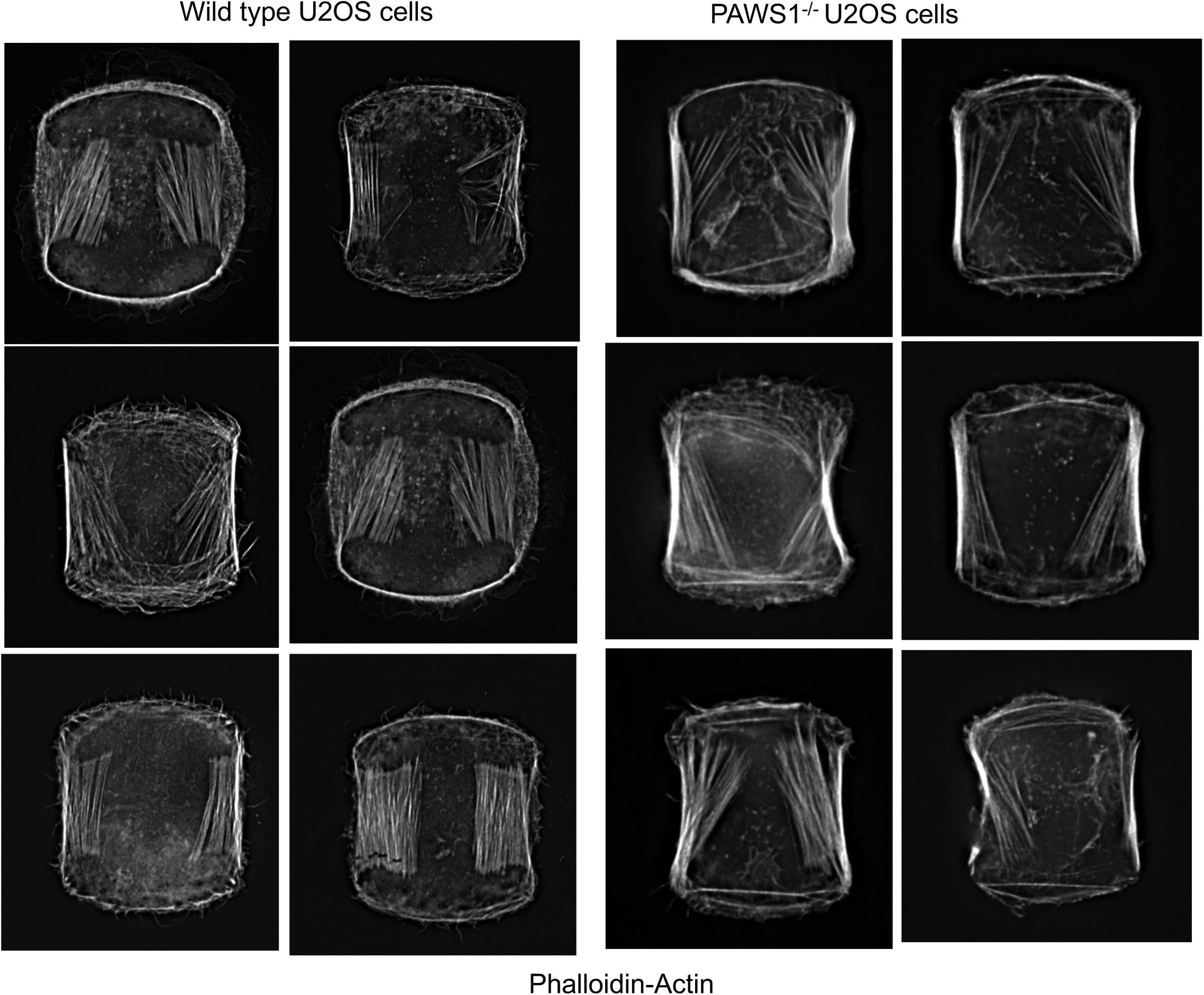
Additional representative images of micropattern adhesion dynamics of phalloidin-stain in wild type and PAWS1^-/-^ U2OS cells in the double crossbow H-micropattern. Monochrome images of each genotype are presented to indicate the shape and structure of actin fibres. Cells and images were processed as described in Legends to Figure 3C.

**Supplementary Figure 6.**
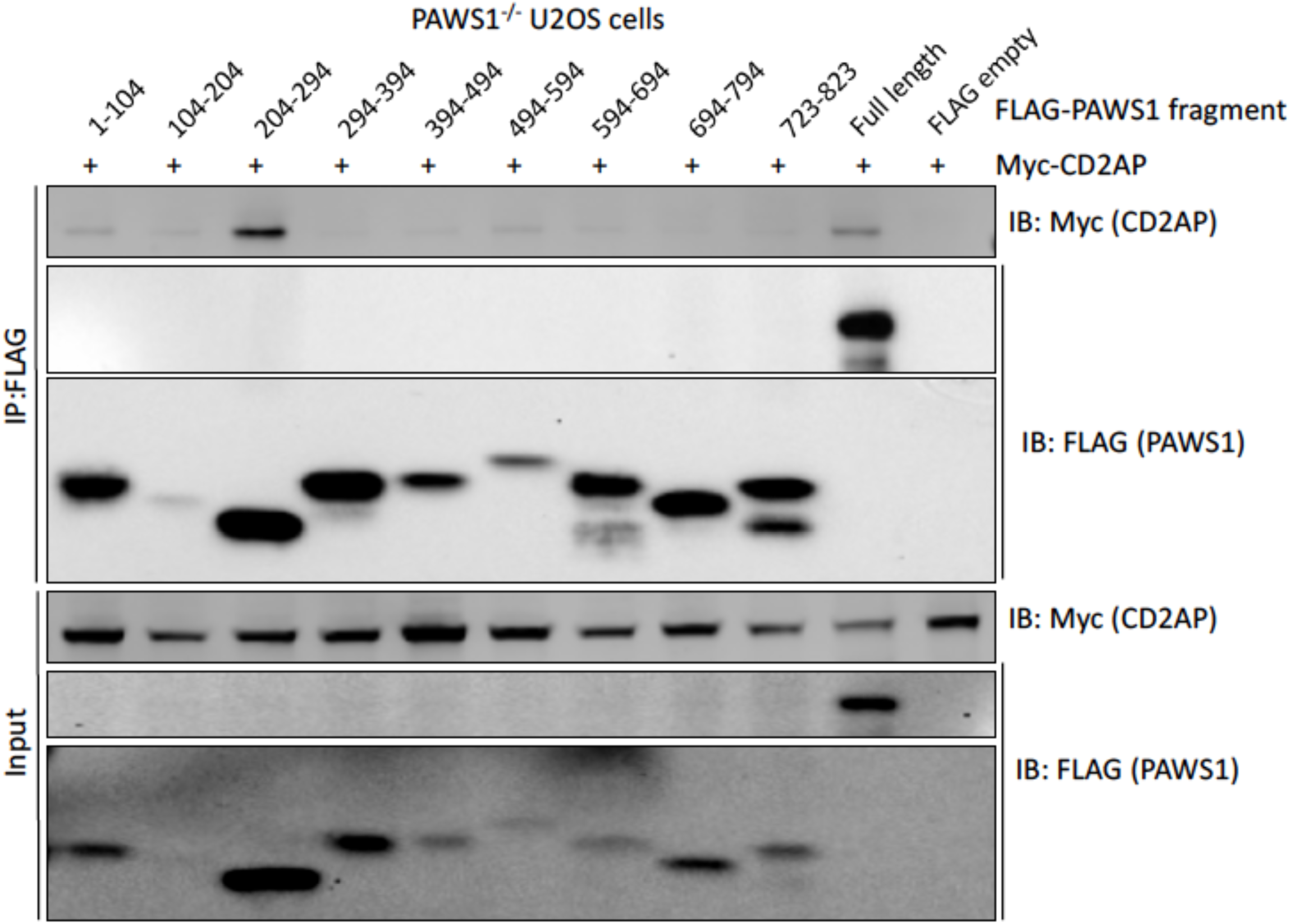
Mapping PAWS1-CD2AP interactions. The indicated fragments of Flag-tagged PAWS1 were co-expressed with Myc-tagged full-length CD2AP in PAWS1^-/-^ U2OS cells for 48 hrs. Extracts or anti-Flag IPs were subjected to immunoblotting with anti-myc (CD2AP) and anti-Flag (PAWS1) antibodies as indicated.

**Movie 1. 2-dimensional lateral migration live-cell imaging using widefield microscopy.** Live cell imaging of wild type U2OS cells on the left and PAWS1^-/-^ U2OS cells on the right. Upon removal of the wound barrier, cells were allowed to migrate onto the gap for 16 hours. Images were acquired at 40X magnification every 5 minutes continuously.

**Movie 2**. Live cell time-lapse fluorescence microscopy video of PAWS1^-/-^ U2OS cells transfected with mApple-LifeAct (actin tracker) and either **A.** GFP or **B.** GFP-PAWS1 to visualize live cell actin dynamics. Images were acquired every 2 minutes over the course of a 25-minute time frame using a 60X magnification. Scale bars are 20 µm.

**Movie 3. Analysis of focal adhesion dynamics in wild type (left panel) and PAWS1^-/-^ (right panel) U2OS cells using TIRF microscopy**. Wild type **(A)** and PAWS1^-/-^ **(B)** U2OS cells were transfected with RFP-zyxin (red) and Emerald-LifeAct (green) to measure changes in focal adhesions over a time course of 25 minutes. Images were acquired continuously every 60 seconds for 25 minutes. Scales bar indicates 20 µm.

**Movie 4.** TIRF microscopy of PAWS1-GFP and mCherry CD2AP in PAWS1^-/-^ U2OS cells. Cells were transfected for 24 hours with PAWS1-GFP (green) and mCherry CD2AP (red) to measure changes in the co-localization distribution over a time course of 25 minutes. **A**. merged; **B.** PAWS1-GFP (green) only and **C.** mCherry CD2AP (red) only. The movement of co-localized some punctate structures highlight the dynamic nature of PAWS1 and CD2AP complexes during the time course. Scale bars are 10 µm.

